# Distinct cellular processes drive motor skill learning in the human brain

**DOI:** 10.1101/2025.09.22.677842

**Authors:** Guillermina Griffa, Marco Palombo, Abraham Yeffal, Hong-Hsi Lee, Susie Y Huang, Valeria Della-Maggiore

**Affiliations:** IFIBIO Houssay, School of Medicine, Department of Physiology, University of Buenos Aires, Argentina; Cardiff University Brain Research Imaging Centre (CUBRIC), School of Psychology and School of Computer Science and Informatics, Cardiff University, Cardiff, UK; Athinoula A. Martinos Center for Biomedical Imaging, Department of Radiology, Massachusetts General Hospital, Harvard Medical School, Boston, MA, USA; ICIFI School of Science and Technology (ECyT), University of San Martin, Buenos Aires, Argentina; Department of Neurology and Neurosurgery, McGill University, Montreal, QC, H3A 2B4, Canada

## Abstract

Despite decades of research, the biological mechanisms by which motor skills consolidate in the human brain remain poorly understood. Diffusion MRI provides a unique opportunity to probe biological processes non-invasively, as water displacements occur on the micrometer scale. Using diffusion tensor imaging (DTI), our team showed that motor sequence learning (MSL) induces microstructural changes in the hippocampus and key motor regions, suggesting that declarative and procedural systems may operate as part of the same network. Yet DTI cannot identify the cellular source of these changes, leaving open whether they reflect structural plasticity —remodeling of dendritic and astrocytic processes described in rodents— or transient homeostatic responses that accompany learning —neuronal and astrocytic swelling. Here, we combined ultra-high-gradient diffusion MRI with the compartment-based Soma and Neurite Density Imaging (SANDI) model to disentangle the cellular basis of motor skill memory consolidation. DTI showed that MSL induced rapid microstructural changes in the hippocampus, precuneus, and motor regions, but only those in the precuneus and posterior parietal cortex (PPC) persisted overnight. SANDI revealed that DTI changes were driven by two distinct cellular processes: a transient enlargement of the cell soma across all regions consistent with a short-lived homeostatic response, and a sustained rise in cell-process density restricted to the precuneus and PPC, compatible with structural plasticity. By decomposing diffusion signals into their cellular sources, our work disambiguates transient and enduring processes, providing the first non-invasive evidence for the cellular basis of human motor memory consolidation and a framework for studying neuroplasticity in vivo.

## INTRODUCTION

Learning triggers persistent modifications in neuronal and astroglial structure, including synaptogenesis, the remodelling of dendritic spines (Lamprecht and LeDoux, 2004; Holtmaat and Svoboda, 2009), and adjacent astroglial processes, necessary for synapse stabilization (Bernardinelli et al., 2014). These lasting changes are collectively referred to as structural plasticity and form the biological basis of memory consolidation. Yet, learning also elicits transient -reversible-homeostatic responses to increased activity, such as neuronal and astrocytic swelling, which may temporarily alter the physical structure of the cell without leading to structural plasticity (Le Bihan et al., 2006; Jin et al., 2013; Abe et al., 2017; Mader et al., 2019). Disentangling these overlapping processes is essential for a mechanistic understanding of memory consolidation in the human brain.

By capturing water displacements on the micrometer scale, diffusion MRI (dMRI) provides a unique non-invasive window into the cellular architecture of the living human brain (Novikov et al., 2014; Novikov et al., 2019). Using the classic diffusion tensor imaging (DTI) model, we have shown that different motor learning paradigms induce hippocampal and cortical microstructural changes, the persistence of which depends on the explicit/implicit nature of the task and the amount of training (Jacobacci et al., 2020a; Deleglise et al., 2023; Della-Maggiore, 2024; Griffa et al., 2025). Our work suggests that during motor memory consolidation, declarative and procedural systems may operate as part of a unified network. Yet, DTI reflects the averaged diffusion signal from all cellular compartments within a voxel and therefore cannot distinguish plastic from non-plastic processes triggered by learning (Blumenfeld-Katzir et al., 2011).

To address this limitation, here we combined ultra-high-gradient diffusion MRI (Huang et al., 2015, 2020, 2021; Jones et al., 2018; Fan et al., 2018, 2020, 2021, 2022; Ramos-Llordén et al., 2025) with the Soma and Neurite Density Imaging (SANDI, Palombo et al., 2020) model to disentangle, for the first time in humans, the cellular processes underlying motor skill memory consolidation. Building on our prior work (Jacobacci et al., 2020a), we applied the same motor sequence learning (MSL) paradigm on the Connectome scanner (300 mT/m; Setsompop et al., 2013; McNab et al., 2013; Jones et al., 2018; Fan et al., 2022) and acquired multi-shell dMRI at baseline, 30 min, and 24 h post-training (Figure 1). SANDI decomposes the diffusion signal into contributions from somas, neurites, and extracellular space, providing biological specificity unattainable with conventional DTI (Palombo et al., 2020; Ianuş et al., 2022; Krijnen et al., 2023; Lee et al., 2024; Krijnen et al., 2024). By applying SANDI to longitudinal dMRI data, we tracked dynamic changes in cell bodies and processes across distinct timescales, thereby disambiguating short-lived homeostatic responses from enduring structural plasticity. Our findings show that distinct cellular processes acting on different timescales support human motor memory consolidation, resolving a long-standing limitation in how diffusion imaging of gray matter is interpreted in the context of plasticity.

**Figure 1.**
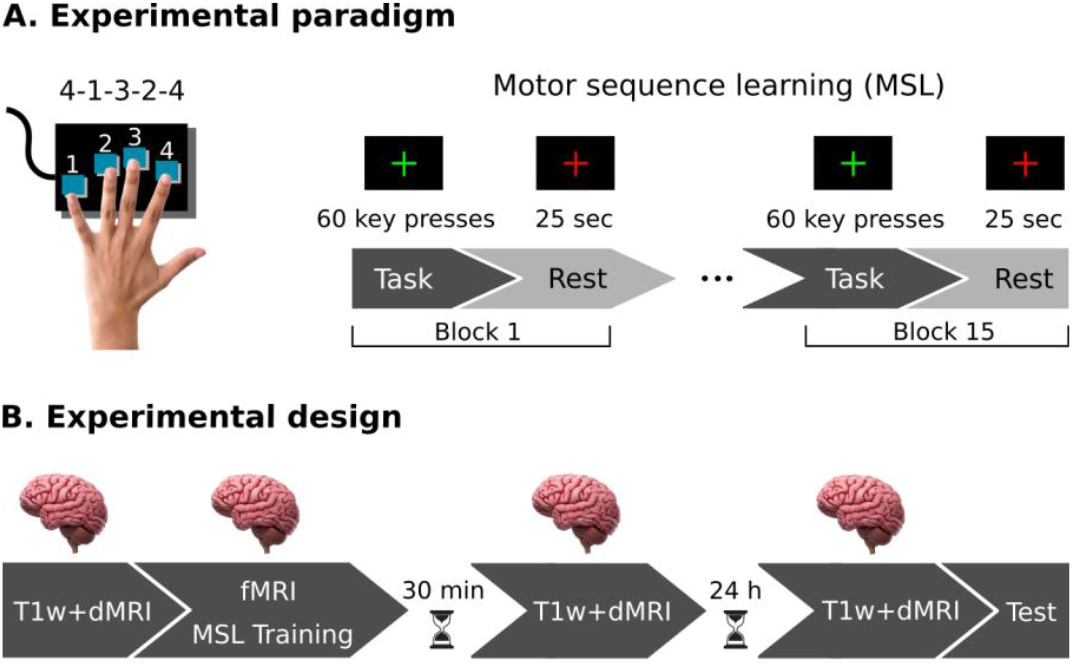
Experimental paradigm and design. **A)** *Experimental paradigm*. Right-handed participants performed a motor sequence learning (MSL) task involving a 5-item sequence of finger movements on a keyboard, using the four fingers of their non-dominant hand (4-1-3-2-4, with 4 representing the index finger and 1 representing the pinky finger). Initially, subjects memorized the sequence and were then instructed to execute it as quickly and accurately as possible during task blocks, and relax during rest periods. A total of 15 blocks were completed. **B)** Experimental design. fMRI images were obtained during the MSL task following a block design. To assess structural plasticity induced by learning, ultra-high multi-shell dMRI and corresponding T1-weighted images (T1w) were acquired longitudinally at three time points: before learning, 30 min post-learning, and 24 h post-learning. Finally, to assess overnight offline gains, participants completed 8 additional practice blocks 24 h after training; the test was assessed after the MRI session.

## RESULTS

Participants learned a 5-item motor sequence using the fingers of their non-dominant hand. As expected, MSL on the Connectome scanner reproduced the behavioral, functional MRI (fMRI), and DTI findings of our previous work conducted on a clinical scanner (Jacobacci et al., 2020a), providing the foundation for the compartment-resolved analyses with SANDI. Specifically, gains in performance in MSL occurred during the rest periods interleaved with practice (micro-offline gains, MOGs), rather than during sequence execution (micro-online gains, MONGs) (Figure 2A). fMRI analysis revealed that activity in the left hippocampus and right precuneus increased during the rest periods, while cortico-cerebellar and cortico-striatal networks were consistently more active during task execution (Figure 2B).

**Figure 2.**
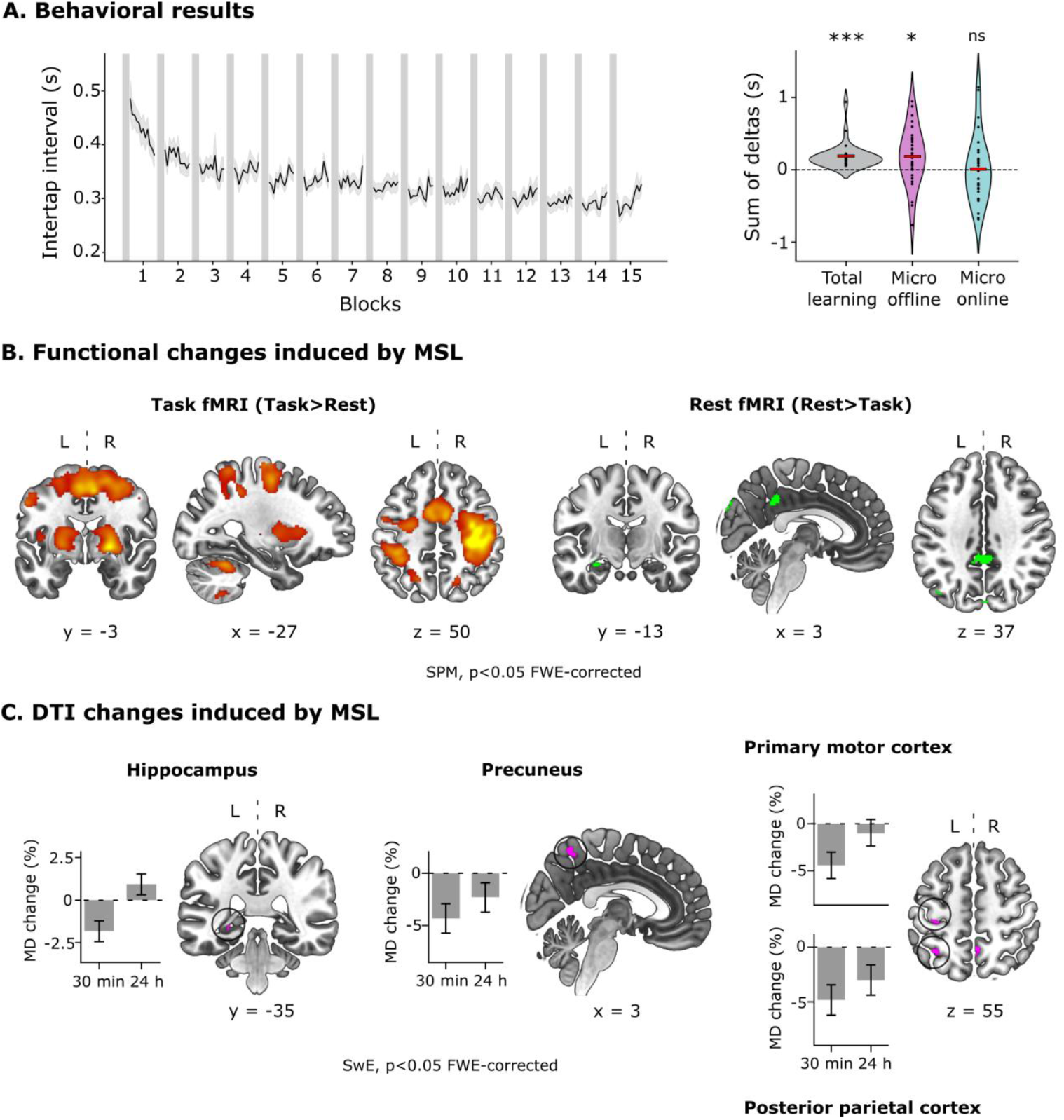
Behavioral, functional, and DTI changes induced by motor sequence learning. **A)** *Behavioral results* - The left panel shows the motor sequence learning curve, depicting the intertap interval -the time elapsed between successive key presses from correctly executed sequences- as a function of blocks (∼12 sequences per block). The right panel shows the metrics corresponding to micro-offline gains (MOGs), micro-online gains (MONGs), and total learning, which were obtained based on the cumulative sum of deltas in performance during practice and rest blocks. Specifically, MOGs represent the difference in the mean intertap interval between the last correct sequence of a block and the first correct sequence of the next block. In contrast, MONGs represent the difference between the first and last correct sequence within a block. Total learning=MONGs+MOGs. Note that, in line with our previous work, improvements in performance on the MSL task occurred during rest periods interleaved with practice (t-test against zero; p=0.034), rather than during sequence execution (p=0.59). Non-significant (ns), ***p<0.001, *p<0.05 corrected by Bonferroni. **B)** *Functional results -* Shown are the whole-brain voxelwise statistical parametric maps (SPMs) for the Task (Task > Rest; left plot) and Rest (Rest > Task; right plot) conditions (p<0.05, Family-Wise Error [FWE]-corrected). Note that cortico-cerebellar and cortico-striatal networks were more active during task execution (in red), whereas the hippocampus and the precuneus were more active during the rest periods (in green). **C)** *DTI results -* Shown are the results from the Sandwich Estimator (SwE) statistical model on mean diffusivity (MD) using a whole-brain threshold-free cluster enhancement (TFCE) approach (p<0.05 FWE-corrected) across the three dMRI sessions (baseline, 30 min, 24 h); barplots represent the mean % change of MD relative to the baseline ± 95% confidence intervals (CI) for the clusters identified in the whole-brain analysis. Note that MD changes are expressed as % of the baseline to facilitate the comparison to our DTI analysis carried out in a clinical MRI scanner (Jacobacci et al., 2020a). MSL induced a rapid (30 min) reduction in MD over the left posterior hippocampus, right precuneus, left primary motor cortex (M1), and left posterior parietal cortex (PCC). Remarkably, while hippocampal and M1 changes were transient, returning to baseline within 24 hours, those in the precuneus and PPC persisted overnight.

Finally, DTI analysis revealed rapid decreases in mean diffusivity (MD) in the left posterior hippocampus and precuneus (Figure 2C), replicating our prior findings. Additional MD reductions were detected in the primary motor and posterior parietal cortex—two brain regions implicated in skill acquisition (Karni et al., 1998; Doyon et al., 2003)—which were likely detected in this study due to the larger sample size (29 vs 20 subjects).

In sum, motor sequence learning on the Connectome scanner reproduced the expected behavioral, fMRI, and DTI results from our previous work, confirming the reliability of this dataset for further analyses.

### Distinct cell processes govern DTI changes in the hippocampus and key motor regions

Visual inspection of MD temporal patterns displayed in Figure 2C points to the existence of two processes driving microstructural changes, potentially shedding light on the mechanisms underlying memory consolidation: a short-lasting process, involving transient -reversible-changes in the microstructure of the left posterior hippocampus and M1, and a long-lasting process impacting on the precuneus and PPC. These were statistically confirmed through two separate whole-brain analyses (Figure 3A and 3B, DTI).

**Figure 3.**
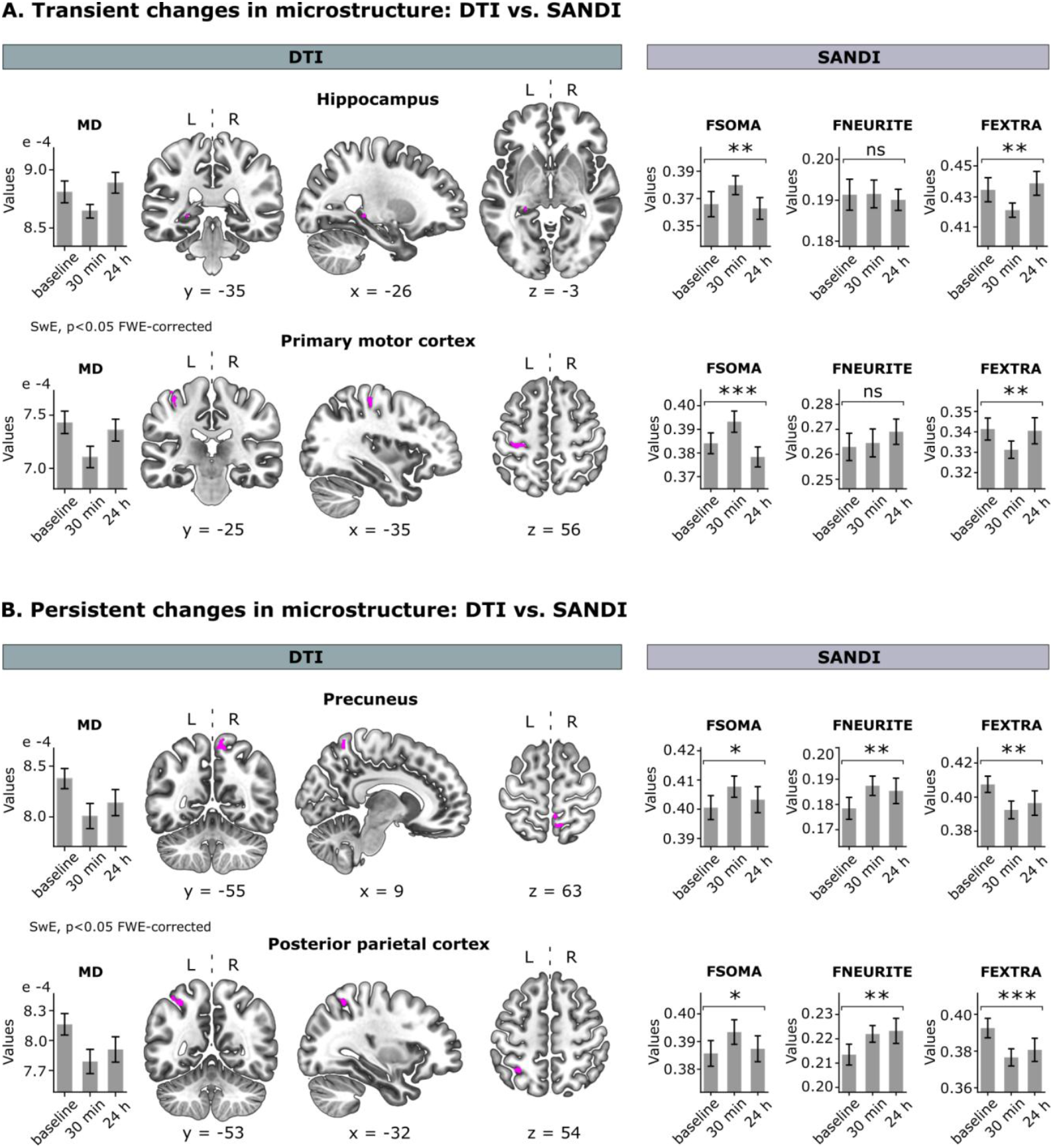
Temporal dynamics of microstructural changes induced by MSL: DTI vs. SANDI. **A)** The MSL task induced a transient decrease in MD (left panel) in the left posterior hippocampus and left primary motor cortex (SwE, p<0.05 FWE-corrected). These changes were accompanied by a transient increase in apparent cell-body density (FSOMA; right panel; repeated-measures ANOVA, Bonferroni-corrected; hippocampus: p = 0.016; M1: p<0.001). **B)** MSL also induced a long-lasting reduction in MD (left panel) in the precuneus and posterior parietal cortex (SwE, p<0.05 FWE-corrected), which was associated with both a transient increase in apparent cell-body density (FSOMA; right panel; precuneus: p=0.044; PPC: p=0.043) and a sustained increase in apparent cell-process density (FNEURITE; right panel; repeated-measures ANOVA, Bonferroni-corrected; precuneus: p = 0.024; PPC: p = 0.0071) that persisted at least 24 h post-learning. Bar plots depict the time course of the mean MD/SANDI metrics ± 95% CI for each brain region. *p<0.05, and **p<0.05; ***p<0.001 Bonferroni-corrected.

To disentangle the cellular compartments contributing to these temporal patterns, we applied the SANDI model to the multi-shell dMRI dataset and carried out an ROI statistical approach within the clusters identified by the DTI analysis (Figures 3A and 3B, SANDI). The recently introduced SANDI model (Palombo et al., 2020) accounts for cellular contributions to gray matter by modeling cell bodies as spheres and neurites as cylinders of negligible caliber (namely “sticks”), producing three additive fractions: FSOMA, tackling the contribution of cell soma to the diffusion signal, FNEURITE estimating the contribution of cell processes, including neurites (e.g, dendrites) and processes from glial cells and FEXTRA, which represents the extracellular fluid. Given that FEXTRA is computed from the other two, we focused on the FSOMA and FNEURITE fractions, which are estimated by the model.

Our findings revealed that rapid decreases in MD were associated with transient increases in apparent cell-body density across all four brain regions detected in the whole-brain analyses (Figures 3A and 3B, SANDI, FSOMA). Note that, given the timeline of this study (24 h), we expected changes in the FSOMA fraction to reflect an increase in the size of cell bodies rather than in their number (Pereira et al., 2007; Deng et al., 2010). The fact that these modifications were transient and impacted all bran regions regardless of their embryonic origin (neocortex vs. archicortex) points to non-specific homeostatic processes typically accompanying synaptic potentiation, such as neuronal and/or astrocytic swelling (Jin et al., 2013; Abe et al., 2017; Mader et al., 2019). Note that although swelling may accompany synaptic plasticity (e.g., to maintain electrolytic balance during LTP), it is a short-lived functional response different from structural remodeling.

In contrast, sustained MD changes went hand in hand with a persistent increment in apparent cell-process density within the PPC and precuneus (Figure 3B, SANDI, FNEURITE). This morphological modulation is consistent with structural plasticity observed in non-human animals involving the remodeling of neuronal and astrocytic processes (Anderson et al., 1994; Holtmaat and Svoboda, 2009; Wilbrecht et al., 2010; Bernardinelli et al., 2014). Critically, while the different timescales of changes can be inferred from the temporal pattern of MD (Figure 2C), MD alone does not provide information about the biological nature of microstructural changes. Our findings suggest that memory consolidation involves both transient and persistent cellular processes, potentially reflecting distinct biological mechanisms.

Interestingly, FEXTRA (Figures 3A and 3B, FEXTRA) showed similar temporal dynamics to MD across all brain regions. This pattern supports the hypothesis proposed by Assaf and colleagues, suggesting that reductions in MD may reflect a decrease in interstitial space due to cellular expansion, such as astrocytic hypertrophy triggered by LTP-like plasticity (Sagi et al., 2012).

### Topographical correspondence of hippocampal changes detected with DTI and SANDI

The results presented above arise from applying the SANDI model to clusters identified through the DTI whole-brain analysis. To evaluate whether the sensitivity to learning-related changes in gray matter microstructure was comparable across models, we fitted DTI and SANDI independently for each hippocampal subfield and contrasted their topographical distributions. For this purpose, we segmented the bilateral hippocampus along its longitudinal axis into head, body, and tail regions (Iglesias et al., 2015), based on each subject’s T1-weighted images.

We found a significant anatomical overlap for MD and FSOMA across the three subfields (Figure 4). Specifically, transient decreases in MD in the tail and body of the left hippocampus were matched by a transient increase in apparent cell-body density. Notably, no significant changes in these metrics were detected in either the head or the right hippocampus. Additionally, MSL did not modulate apparent cell-process density in any subfield. Collectively, these results are consistent with the output from the whole-brain analysis, converging on the left posterior hippocampus.

**Figure 4.**
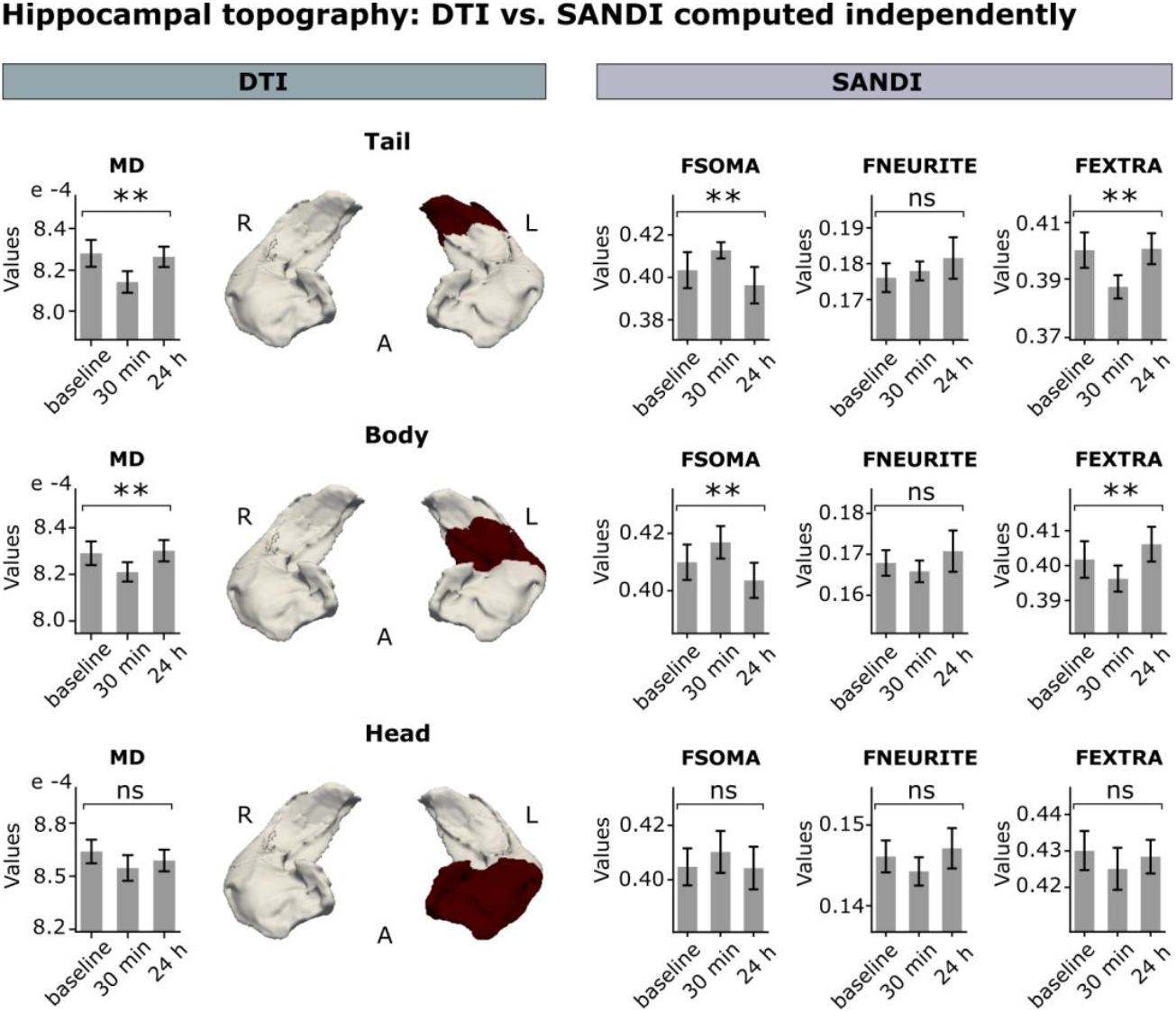
MSL-induced microstructural changes assessed independently using DTI and SANDI. Both MD and SANDI metrics were computed separately across the tail, body, and head subfields of the hippocampi (highlighted in red), which were segmented based on each subject’s T1-weighted images (Iglesias et al., 2015). We observed anatomical overlap for MD and FSOMA metrics across the body and tail of the left hippocampus (repeated-measures ANOVA, Bonferroni-corrected; tail: MD: p=0.0034; FSOMA: p=0.022; body: MD: p=0.034; FSOMA: p=0.026). Notably, no changes were identified by either model in the hippocampal head or in the right hippocampus. MSL did not affect apparent cell-process density in any of the subfields (p>0.05). Bar plots depict the time course of mean MD/SANDI metrics ± 95% confidence intervals (CIs). **p<0.05; ns = non-significant.

The localization of microstructural changes to the posterior region of the left hippocampus aligns with findings from our previous MSL work (Jacobacci et al., 2020a) and a recent study examining the hippocampus’s involvement in visuomotor adaptation—a form of motor learning related to skill recalibration (Griffa et al., 2025). The similar topography across DTI and SANDI reinforces the reliability of both models in detecting microstructural changes and affirms the robustness of the SANDI model.

## DISCUSSION

Tracking the emergence of neuroplasticity is not only essential for understanding memory encoding and consolidation but also for identifying early markers of brain dysfunction, such as neuroinflammation (Ligneul et al., 2019; Lee et al., 2021; Genovese et al., 2021; De Marco et al., 2022; Garcia-Hernandez et al., 2022). In this study, we laid the foundation for inferring structural plasticity non-invasively in humans by combining high-gradient diffusion MRI with advanced modeling techniques. We explored the temporal dynamics of cell bodies and their processes using SANDI, a compartment-based approach modeling diffusion at the sub-voxel level (Palombo et al., 2020), and compared it to the traditional DTI method (Basser et al., 1994; Sagi et al., 2012). While both models revealed similar patterns, key differences emerged in the temporal dynamics of microstructural changes. Both MD and apparent cell-soma density showed transient changes shortly after learning, with a decrease in MD aligning with an increase in soma fraction in the hippocampus and key motor regions critical for MSL acquisition. However, MD reductions persisting overnight were linked to a sustained increase in apparent cell-process density in the same brain regions. SANDI thus uncovered distinct short and long-lasting cellular processes that were undetectable using DTI alone.

In the last decade, many multi-compartment mathematical models have been proposed to overcome the limitations of DTI in mapping brain tissue microstructure in the living human brain (Alexander et al., 2010; Zhang et al., 2012; Kaden et al., 2016; Novikov et al., 2018; Alexander et al., 2019; Fan et al., 2020; Huang et al., 2020). However, they are based on geometrical assumptions specific to white matter and are not optimized to study plasticity in gray matter. By providing deeper insights into the cellular compartments modulated by learning, SANDI offers clues about potential mechanisms involved in motor memory consolidation. We hypothesize that the transient increase in apparent cell-soma density observed shortly after learning may reflect non-specific homeostatic processes accompanying synaptic plasticity, such as neuronal and astrocytic swelling triggered by neuronal activity (Le Bihan et al., 2006; Sagi et al., 2012; Jin et al., 2013; Abe et al., 2017; Mader et al., 2019). In contrast, the persistent increase in the apparent cell-processes density likely reflects structural plasticity in the form of remodeling (Anderson et al., 1994; Kleim et al., 2007; Holtmaat and Svoboda, 2009; Wilbrecht et al., 2010; Stevenson et al., 2021). Future research combining a longitudinal design with the estimation of cell membrane permeability, currently being implemented by our team based on multiple-diffusion-time dMRI acquisition (Lee et al., 2022; Jelescu et al., 2022; Olesen et al., 2022; Chan et al., 2024) would help address this hypothesis. Our mechanistic approach thus constitutes a critical first step forward in distinguishing between microstructural changes driven by plastic versus non-plastic processes. This distinction is critical since, until now, the lack of anatomical specificity of dMRI models forced scientists to rely on indirect metrics that, while sensitive to gray matter microstructure, may not necessarily reflect plasticity itself (Hofstetter et al., 2017; Hofstetter and Assaf, 2017; Jacobacci et al., 2020a; Griffa et al., 2025).

The observed short- and long-term changes in the cell body and neurite compartments were likely functionally linked, as both may have been modulated either indirectly (e.g., through homeostatic swelling) or directly (e.g., through the remodeling of neuronal and glial processes) during memory consolidation. However, the fact that only transient changes in apparent cell-body density were detected in the hippocampus and M1 raises the question of whether these brain regions are involved in MSL consolidation at all. Recent work from non-human animals (Goto et al., 2021) suggests that there are several waves of synaptic plasticity, with the first one acting locally within the hippocampus for context specificity and the second in the neocortex during sleep (known as systems consolidation). Within this framework, it is possible that hippocampal changes are transient because they do not persist through systems consolidation but persist in the precuneus and parietal cortex as they form part of the extended network involved in motor memory storage. Alternatively, given that procedural memories require repeated practice for reinforcement (Wymbs et al., 2012, 2015), the transient nature of changes in hippocampal microstructure may reflect the limited amount of training (15-25 minutes) to which volunteers were exposed. Indeed, recent evidence from our team suggests that motor tasks with a strong implicit learning component or extended training over multiple visuomotor adaptation sessions lead to persistent reductions in MD (Griffa et al., 2025). This raises the possibility that further practice might indeed lead to structural plasticity in the hippocampus, a hypothesis amenable to testing.

In line with previous work from our team, changes in hippocampal microstructure were localized to the posterior portion of the left hippocampus, specifically over the body and tail (Jacobacci et al., 2020a; Deleglise et al., 2023; Griffa et al., 2025). This left lateralization, which mirrors that observed in M1 and PPC—two key regions for MSL acquisition (Karni et al., 1998; Doyon et al., 2003)—may originate from the asymmetric control of the motor system during the planning of skilled and reaching movements (De Renzi and Lucchelli, 1988; Schaefer et al., 2007; Yang et al., 2024; Merrick et al., 2022). Anatomical evidence indicates that the anterior and posterior hippocampus indeed form part of two distinct networks associated with processing the “what” and “where” aspects of stimuli, respectively (Rolls et al., 2023). Specifically, the posterior portion of the hippocampus ultimately connects with the PPC involved in motor planning and MSL through the retrosplenial cortex, and the precuneus identified herein (Della-Maggiore et al., 2004; Kahn et al., 2008; Margulies et al., 2009; Vesia et al., 2010; Adnan et al., 2016; Dalton et al., 2022). Therefore, we propose that the consistent topography of hippocampal changes observed with both DTI and SANDI, and recently replicated by our team in other motor paradigms (Griffa et al., 2025), may be attributed to the connectivity between the limbic and motor systems.

Despite these advances, some potential limitations should be considered. While the magnitude of the cell-body and cell-process density estimated by SANDI is in line with those reported in other human studies (Lee et al., 2024; Karat et al., 2024), the cell-process density was generally lower than values observed in histology, a well-known limitation of dMRI models (Lampinen et al., 2023). This discrepancy may stem from the assumption in SANDI of no water exchange between compartments, which may have been partially violated even at the short diffusion times used on the Connectome scanner (24 ms) (Jelescu et al., 2022). Nevertheless, this limitation is unlikely to impact on the validity of our findings, as the learning-related changes reported here were interpreted relative to the baseline for each brain region. Another consideration is that SANDI estimates signal fractions rather than absolute proton densities, which reflect the concentration of hydrogen protons—and, by extension, water content—in each compartment. As a result, the estimated contributions of the soma, neurites, and extracellular space are interdependent. More accurate, compartment-specific quantification would require diffusion-relaxometry acquisitions with longer scan times (Afzali et al., 2021; Gong et al., 2023), calibrated against a proton density reference such as cerebrospinal fluid (Mezer et al., 2016). Lastly, while this study lacked a dedicated control condition that did not learn, it replicated our previous findings in which learning-related changes in MD were observed relative to a control involving non-sequential finger tapping (Jacobacci et al., 2020a). The successful replication of both our functional and DTI results on the Connectome MRI scanner strengthens confidence in the specificity of our findings.

In conclusion, our study provides new insights into the cellular mechanisms underlying motor skill learning in humans. By combining ultra-high-gradient diffusion MRI with the SANDI model, we successfully dissociated transient and persistent changes in cell bodies and cell processes within the hippocampus and motor regions, likely reflecting distinct biological mechanisms involved in memory consolidation. Tracking these processes non-invasively across timescales represents a significant advance, as previous gray matter models could not distinguish plastic from non-plastic structural change. By resolving temporal dynamics at the cellular level, our approach establishes a mechanistic framework to study neuroplasticity in vivo and provides a crucial bridge between experimental neuroscience in animals and human brain imaging. Future work exploiting this framework — for example, incorporating multiple diffusion times — holds promise for advancing our understanding of neuroplasticity in both health and disease.

## METHODS

### Participants

29 healthy human subjects between 18 and 36 years old (16 female, mean age ± SD = 26.86 ± 5.41 years) were recruited under approval by the Institutional Review Board of Mass General Brigham. Written informed consent was obtained from all participants. All participants reported no psychiatric, neurological, or cognitive impairment, nor any history of sleep disturbances. All subjects were right-handed as assessed by the Edinburgh Handedness Inventory (Oldfield, 1971).

### Experimental paradigm

Both the learning paradigm and the experimental design used in this study were identical to our previous MSL study (Jacobacci et al., 2020a). Participants were instructed to learn a 5-item motor sequence using the four fingers of their non-dominant (left) hand (4-1-3-2-4, 4 being the index finger and 1 being the pinky finger) as quickly and accurately as possible (Figure 1A).

### Experimental design and procedure

All participants performed the MSL task in a self-paced manner during 15 blocks of 12 sequences each (60 key presses) interleaved with rest periods of 25 s (Jacobacci et al., 2020a). The task lasted around 15-20 minutes. Multi-shell Diffusion MRI, T1-weighted, and functional images were acquired throughout the study. fMRI was collected during the MSL task following a block design (task, rest), while dMRI and T1w images were obtained in a longitudinal design at three time points: before practice (baseline), 30 min, and 24 h after learning to track the dynamics of structural plasticity in the short- and long-term (Figure 1B). Finally, to assess overnight offline gains, participants completed 8 additional practice blocks 24 h after training; the test was assessed after the MRI session.

### MRI acquisition

Diffusion MRI, T1-weighted, and fMRI images were acquired on the original 3T Connectome MRI scanner (MAGNETOM CONNECTOM, Siemens Healthcare, Forchheim, Germany), equipped with a maximum gradient strength of 300 mT/m and a slew rate of 200 T/m/s, using a 64-channel phased array head coil (Keil et al., 2013).

Functional images (BOLD) were obtained following a block design during MSL (15 blocks of self-paced practice alternated with 25 s rest). The acquisition parameters were chosen to match our previous MSL study (Jacobacci et al., 2020a): voxel size = 3×3×3 mm3; FOV = 210 mm; 42 slices aligned with the AC-PC line; 10% gap; posterior-anterior (P-A) phase encoding direction; repetition time (TR) = 1440 ms; echo time (TE) = 30 ms; multiband acceleration factor = 2; no PAT; bandwidth (BW) = 1786 Hz/Px; echo spacing = 0.62 ms; EPI factor = 70; flip angle = 69°. A gradient echo (GRE) fieldmap was also acquired for correction of field inhomogeneities using the following parameters: 42 slices; 10% gap; voxel size = 3×3×3 mm3; field-of-view (FOV) = 210 mm; posterior-anterior (P-A) phase-encoding direction; TR = 716 ms; TE1 = 4.92 ms; TE2 = 7.38 ms; flip angle = 55°; BW = 600 Hz/px.

T1w and dMRI images were acquired at baseline, 30 min, and 24 h after training. T1w images were obtained using the multi-echo sequence and the following parameters: TR = 2530 ms; TE = 1.15 ms; flip angle = 7°; Inversion Time (TI) = 1100 ms; BW = 649 Hz/Px; FOV = 210 mm; voxel size = 1×1×1 mm3; parallel acquisition = GRAPPA mode, acceleration factor = 3. The acquisition was performed in sagittal slices. The T1w images were used for the coregistration and subsequent normalization of fMRI images and for the segmentation of each subject’s hippocampal subfields.

Multi-shell dMRI images were acquired using a monopolar pulsed-gradient spin-echo, single-shot echo-planar-imaging sequence (Tian et al., 2022) using the following parameters: diffusion time = 24 ms; gradient pulse duration = 8 ms; TR = 4200 ms; TE = 55 ms; voxel size = 2×2×2 mm3; partial Fourier factor = 6/8, GRAPPA acceleration factor = 2; multiband acceleration factor (MB) = 2; anterior-posterior (A-P) phase-encoding direction. dMRI images were acquired in 8 b-values: 32 gradient directions for b-values=200, 500, 1200, 2400 s/mm2, 64 directions for b-values=4000, 6000, 8000 s/mm2, and 22 interspersed non-diffusion-weighted b = 0 images. The latter were acquired for signal normalization and motion correction, of which five were acquired with reversed phase-encoding direction (posterior-to-anterior) for susceptibility distortion correction.

### MRI preprocessing

#### Functional MRI

Preprocessing of functional images was conducted primarily using Statistical Parametric Mapping (SPM12, Wellcome Department of Cognitive Neurology). Initial corrections addressed gradient non-linearity (Jovicich et al., 2006). The functional time series was motion-corrected using the middle volume of the series as a reference. Since the experimental paradigm consisted of a block design, we did not perform slice-timing correction. Susceptibility distortion due to magnetic field inhomogeneities was corrected using the phase and magnitude images of a gradient-echo field map, and the realignment was performed to the middle volume of each functional time series with the aid of FSL’s “fslsplit” and “fslmerge” commands. T1w structural images were coregistered to the BOLD images and normalized to the standard MNI152 template. This transformation was subsequently applied to the BOLD data, and the functional images were smoothed using a Gaussian kernel with an 8-mm full-width at half-maximum.

#### T1

Human hippocampal subfields were segmented using FreeSurfer (version 7.0). Each subject’s T1w image was processed through FreeSurfer’s ‘recon-all’ pipeline. The hippocampal head, body, and tail were then extracted along the longitudinal axis using the optimized ‘segmentHA_T1’ function (Iglesias et al., 2015).

#### Diffusion MRI

Preprocessing and normalization of the diffusion MRI images were performed using the FMRIB Software Library (FSL; University of Oxford) (version 6.0.3), MRtrix3 (version 3.0.3), and Advanced Normalization Tools (ANTs; Wellcome Department, UCL) (version 2.4.2).

Preprocessing steps were applied separately for each scanning session (baseline, 30 min, and 24 h post-training). Images were denoised using MRtrix3’s “dwidenoise”, corrected for Gibbs ringing using MRtrix3’s “mrdegibbs”, and corrected for the nonlinearity of gradients (Jovicich et al., 2006). Geometric susceptibility distortion correction was made using FSL’s “Topup”. Subsequently, head motion, eddy currents, and b-vector rotation were made using FSL’s “eddy” (Andersson et al., 2016). Finally, bias correction was then applied using MRtrix3’s “dwibiascorrect”.

Following preprocessing, FSL’s “dtifit” was used to fit the diffusion tensor model, generating scalar maps for fractional anisotropy (FA) and mean diffusivity (MD) metrics. FA and MD maps were calculated for each subject and each voxel using the 1200 s/mm2 b-shell. Additionally, the Soma and Neurite Density Imaging (SANDI) model was fitted to the multi-shell dMRI acquisition using the SANDI MATLAB toolbox (https://github.com/palombom/SANDI-Matlab-Toolbox) (Palombo et al., 2020). SANDI accounts for cellular contributions to gray matter microstructure by modeling cell bodies as spheres, neurites as sticks, and extracellular space as isotropic Gaussian diffusion, producing three additive fractions. FSOMA tackles the contribution of the cell body to the diffusion signal; FNEURITE estimates the contribution of cell processes such as neurites and cell processes from glial cells; FEXTRA, which represents the extracellular fluid, is calculated based on the other two fractions (FSOMA + FNEURITE + FEXTRA = 1).

To quantify the DTI and SANDI metrics for each hippocampal subfield in native space (Region of Interest –ROI analysis), MD maps and SANDI-derived fractions were registered to each subject’s T1w image using a linear registration algorithm from ANTs. On the other hand, for the whole-brain analysis, DTI and SANDI maps were normalized to stereotaxic space (MNI152) using a custom-made pipeline based on ANTs to minimize across-session test-retest reproducibility error (Jacobacci et al., 2020b). Finally, the normalized DTI and SANDI maps were smoothed using FSL’s smoothing function with a 4-mm full-width at half maximum Gaussian kernel.

### Data analysis and statistics

#### Behavior

Motor sequence learning was quantified using the intertap interval, defined as the time elapsed between successive key presses during correctly executed sequences (Jacobacci et al., 2020a). The mean intertap interval was calculated for each correct sequence (∼12 sequences per block) and each subject. Then, the intertap intervals from correct sequences were averaged across all subjects for each block.

Gains in performance during MSL were assessed as in our previous study (Jacobacci et al., 2020a) into micro-online gains (MONGs), micro-offline gains (MOGs), and total learning. MONGs were calculated as the difference (delta) between the mean intertap interval of the first and last correct sequence within a practice block, while MOGs were computed as the difference between the mean intertap interval of the last correct sequence of a practice block and the first correct sequence of the following block. Total learning represented the sum of MONGs and MOGs.

To statistically assess gains in performance for each of these behavioral metrics, we performed t-tests against zero, with p-values corrected for multiple comparisons using Bonferroni adjustments.

#### Functional MRI

Statistical analyses of BOLD images were conducted using SPM12. We performed a standard whole-brain voxelwise general linear model (GLM) analysis to identify brain regions more active during motor execution (Task > Rest) and during rest periods (Rest > Task). To determine whether the parameter estimates (beta weights) from this GLM were significantly different from zero, we applied a one-tailed t-test for each condition (p<0.05 corrected for family-wise error–FWE). One subject was excluded from the functional analyses due to a technical inconvenience during acquisition affecting most of the dataset.

#### Diffusion MRI

To assess overall changes in MD across sessions (baseline, 30 min, and 24 h after learning), we conducted a whole-brain longitudinal analysis using a non-parametric permutation-based F test, using the threshold-free cluster enhancement (TFCE) approach (H=2, E=0.15) (Smith and Nichols, 2009) implemented with the Sandwich Estimator (SwE) toolbox (version 2.0.0). SwE was specifically designed for accurate modeling of longitudinal and repeated-measures neuroimaging data (Guillaume et al., 2014). The SwE analysis was conducted based on 1,000 permutations, and significant clusters were identified using an FWE-corrected p-value < 0.05. To reveal the temporal dynamics for each cluster of the F test, we extracted the median MD for each cluster, subject and session, expressed it as the percent change relative to the baseline (Figure 2C) and determined the 95% confidence intervals using the summarySEwithin function from the Rmisc package in R (Morey et al., 2008; Team, R. C, 2020). Note that mean % MD changes are shown in Figure 2C to facilitate the comparison between our DTI analysis carried out in a clinical MRI scanner (Jacobacci et al., 2020a) and that acquired at the Connectome I scanner.

To distinguish between short- and long-term changes in microstructure, we conducted two additional whole-brain analyses on the same MD maps using the same SwE approach described above. We then extracted SANDI metrics (FEXTRA, FNEURITE, and FSOMA fractions) from the MD clusters, which were used as ROIs to assess longitudinal metrics of these changes (repeated-measures ANOVA; p<0.05, corrected by Bonferroni). To reveal the temporal dynamics for SANDI metrics, we then computed each fraction’s median for each cluster, subject, and session and the corresponding 95% CI (Figure 3A and 3B).

To contrast DTI and SANDI across hippocampal subfields (Figure 4), we carried out independent statistical analyses (repeated-measures ANOVA; p < 0.05, corrected by Bonferroni) for each metric in the tail, body, and head hippocampi used as ROIs (Iglesias et al., 2015). One subject was excluded from the statistical analysis due to missing data at the 24 h post-training MRI session.

## ACKNOWLEDGMENTS

This research was supported by the Argentinian Agency for the Promotion of Science and Technology (FONCyT) under award numbers PICT 2018-1150 and PICT 2019-2156, the Office of The Director (OD) of the National Institutes of Health (NIH) in partnership with the National Institute of Dental & Craniofacial Research (NIDCR) under award number DP5 OD031854, the National Institute of Aging of the NIH under award number R21 AG085795, the National Institute of Neurological Disorders and Stroke of the NIH under award number R01 NS118187, and the National Institute of Biomedical Imaging and Bioengineering (NIBIB) of the NIH under award numbers U01 EB026996, P41 EB015896, and P41 EB030006.

## Author contributions

V.D-M. conceived the study; V.D-M., M.P., and S.Y.H. designed the research; G.G. performed the experiments; G.G. and A.Y. analyzed the data; M.P. and H-H.L. contributed unpublished analytic tools; G.G., M.P., and V.D-M. interpreted the results; G.G. and V.D-M. wrote the manuscript, while M.P.. H-H.L. and S.Y.H. edited it; V.D-M. and S.Y.H. supervised the project. All authors reviewed and approved the final manuscript.

